# Spatial coordination of peptidoglycan synthesis by MreB directs morphogenesis in *Xanthomonas citri*

**DOI:** 10.64898/2026.11.18.767544

**Authors:** Yue Peng, Thijs de Vroet, Camila Malvessi Pereira, Henrique Ferreira, Dirk-Jan Scheffers

**Author notes:** Correspondence: Dirk-Jan Scheffers.

## Abstract

The cytoskeletal proteins MreB and FtsZ coordinate cell wall synthesis during elongation and division in rod shaped bacteria. How these two systems interact in the citrus canker pathogen *Xanthomonas citri* subsp. *citri* (*X. citri*) remains unclear. Previously, we showed that disruption of FtsZ ring placement, by deletion of *minC*, uncouples cell wall synthesis from division, leading to branching and filamentation. Now, using a MreB-GFP sandwich fusion, we show that MreB localizes at the inner membrane, independently of MinC. Peptidoglycan (PG) synthesis (visualized with the fluorescent D-amino acid analogue HADA) happens mostly at the division septum. Perturbation of MreB with A22 disrupts rod shape maintenance and leads to a dose-dependent growth inhibition. A22 induces a reversible relocalization of MreB into the cell cap region, which is accompanied by a shift of PG synthesis from midcell to the poles. This indicates that MreB is required for proper spatial regulation of PG insertion. This loss of spatial control is exacerbated in *ΔminC* cells, which display increased A22 sensitivity, indicating a synthetic dependence on MreB when FtsZ ring placement is affected. Furthermore, MreB inhibition disrupts ParABS-mediated chromosome segregation, leading to anucleate cells. Our findings establish MreB as a key coordinator of cell wall synthesis, cell shape, and chromosome segregation in *X. citri*

## 1 Introduction

Citrus canker is one of the most common diseases of citrus trees, and is caused by the Gram-negative bacterial species *Xanthomonas citri* subsp. *citri* (*X. citri*) (Ali et al., 2023). *X. citri* is a rod-shaped, aerobic bacterium, with a size of around 1.5-2.0 × 0.5-0.75 μm (Ference et al., 2018). *X. citri* colonies are yellow because the bacteria produce a yellow pigment, xanthomonadin (Cao et al., 2020), which is believed to protect them against solar irradiation ((Rajagopal et al., 1997)).The bacteria can be spread from tree to tree by wind and rain, and enter the plant tissues through stomata and wounds. This results in round yellow or brown patches on the aerial organs of citrus, called canker, which lead to defoliation, reduction of the photosynthetic rate, and premature fruit drop and severe economic harm to citrus producers (Lanza et al., 2019; Liu et al., 2024).

In rod-shaped bacteria, peptidoglycan (PG) synthesis is essential for cell division and elongation, and is regulated by two bacterial protein complexes: the divisome and the elongasome (Szwedziak and Löwe, 2013). FtsZ, the bacterial homologue of tubulin, organizes the divisome. It polymerizes into a ring-like structure, which defines the division site and directs PG synthesis to drive constriction (Barrows and Goley, 2021; Cameron and Margolin, 2024). The localization of the Z-ring is mediated by three Min proteins: MinC, MinD and MinE. As the inhibitor of FtsZ, MinC is activated by MinD to prevent Z-ring assembly at the poles (Ghosal et al., 2014). Topological-specificity factor MinE prevents long term accumulation of MinC/D at midcell, which makes midcell the only available place for Z-ring assembly (Rothfield et al., 2005). In the organization of the elongasome, the bacterial actin homolog MreB plays a critical role. Facilitated by scaffolding and membrane-anchoring proteins such as MreC, MreD, and RodZ, MreB polymerizes into filaments that dynamically associate with the inner membrane, perpendicular to the long axis of the cell. (Typas et al., 2010). This structure interacts with a complex formed by RodA and a dedicated class B Penicillin Binding Protein (bPBP) in diverse non-spherical bacteria. Together, they perform the glycosyl transferase and transpeptidation reactions required to elongate and cross-link the building block LipidII into the growing PG matrix. The orientation and regulation of PG synthesis along the lateral cell wall ensures cylindrical growth of the rod-shape bacteria (Domínguez-Escobar et al., 2011; Grinnell et al., 2022). MreB filament turnover is directly related to PG synthesis (White and Gober, 2012) and controlled by ATP hydrolysis.

In addition to cell elongation and division, bacteria must also complete the replication and segregation of their genomes during each cell cycle (Böhm et al., 2020). The ParABS system organizes chromosome segregation in several bacterial groups including proteobacteria (with some notable exceptions like *Escherichia coli*). ParABS consists of one or more *parS* sites, typically positioned near the origin of chromosome replication (*oriC*) (Livny et al., 2007; Tišma et al., 2024), onto which the DNA-binding protein ParB loads and assembles into a network of protein–DNA complexes (Bock et al., 2022; Funnell, 2016). These complexes interact with the ATPase ParA, generating a ParA–ATP gradient that drives the directional movement of chromosomes toward the cell poles (Jalal and Le, 2020; Vecchiarelli et al., 2010). In various organisms, such as *Caulobacter crescentus* and *E. coli*, deletion or inhibition of MreB has been reported to disrupt chromosome segregation resulting in anucleate cells (Gitai et al., 2005; Iwai et al., 2002), although a direct role for MreB in chromosome segregation has been contested (Karczmarek et al., 2007).

The precise temporal and spatial coordination between chromosome segregation, cell elongation, and division is crucial for the faithful production of daughter cells (Bottomley et al., 2017; Parham et al., 2025). We previously reported that MinC oscillates between cell poles in *X. citri* (Lorenzoni et al., 2017), similar to the oscillation pattern observed in *E.coli* (Wettmann and Kruse, 2018). Deletion of *minC* in *X. citri* results in a characteristic *ΔminC* phenotype, visible by defects in cell division such as the formation of elongated cells, branching cells and minicells. The absence of *minC* also disrupts the spatial overlap localization between PG synthesis and division sites (Lorenzoni et al., 2017). This raised the question whether the delocalized PG synthesis sites would colocalize with MreB and thus, the elongasome. We hypothesize that MreB/the elongasome adopt a more critical role in maintaining cellular integrity when cell division and morphogenesis are disrupted by the absence of MinC. So far, MreB in proteobacteria has been primarily studied in the model organisms *Caulobacter crescentus* (an α-proteobacterium) and *E. coli* (a γ-proteobacterium) (Figge et al., 2004; Takacs et al., 2010). By studying *X. citri* we add a novel γ-proteobacterium to this set, which is sufficiently different from the well-studied models (for example, by containing a *parABS* system, unlike *E. coli*).

In this study, we set out to investigate the role of MreB in cell shape maintenance and PG synthesis. We analyzed how MreB function and localization are impacted by the loss of MinC-mediated division site regulation, and what happens if MreB function is chemically inhibited. The work shows that MreB is essential for the rod-shaped growth of *X. citri*, and our findings provide insight into the functional relationship and coordination between the elongasome and divisome in a γ-proteobacterial plant pathogen.

## 2 Materials and Methods

### 2.1 Bacterial Strains and growth conditions

All strains and plasmids are listed in Table 1. *X. citri* cells were grown at 25℃ in nutrient yeast glycerol broth (NYGB, 5 g/L peptone, 3 g/L yeast extract; 20 g/L glycerol, 15 g/L agar (for solid medium) or in Xam1 medium (2.46 g/L glycerol, 0.247g/L MgSO_4_.7H_2_O, 1.0 g/L (NH_4_)_2_SO_4_, 4.5 g/L KH_2_PO_4_, 10.5 g/L K_2_HPO_4_, 0.5 g/L Na_3_C_6_H_5_O_7_.2H_2_O, 0.3 g/L casamino acids, 1 g/L BSA, pH 5.4 with HCl) (Lorenzoni et al., 2017). MreB-GFP expression was induced by adding 0.05% w/v arabinose.

**Table 1.**
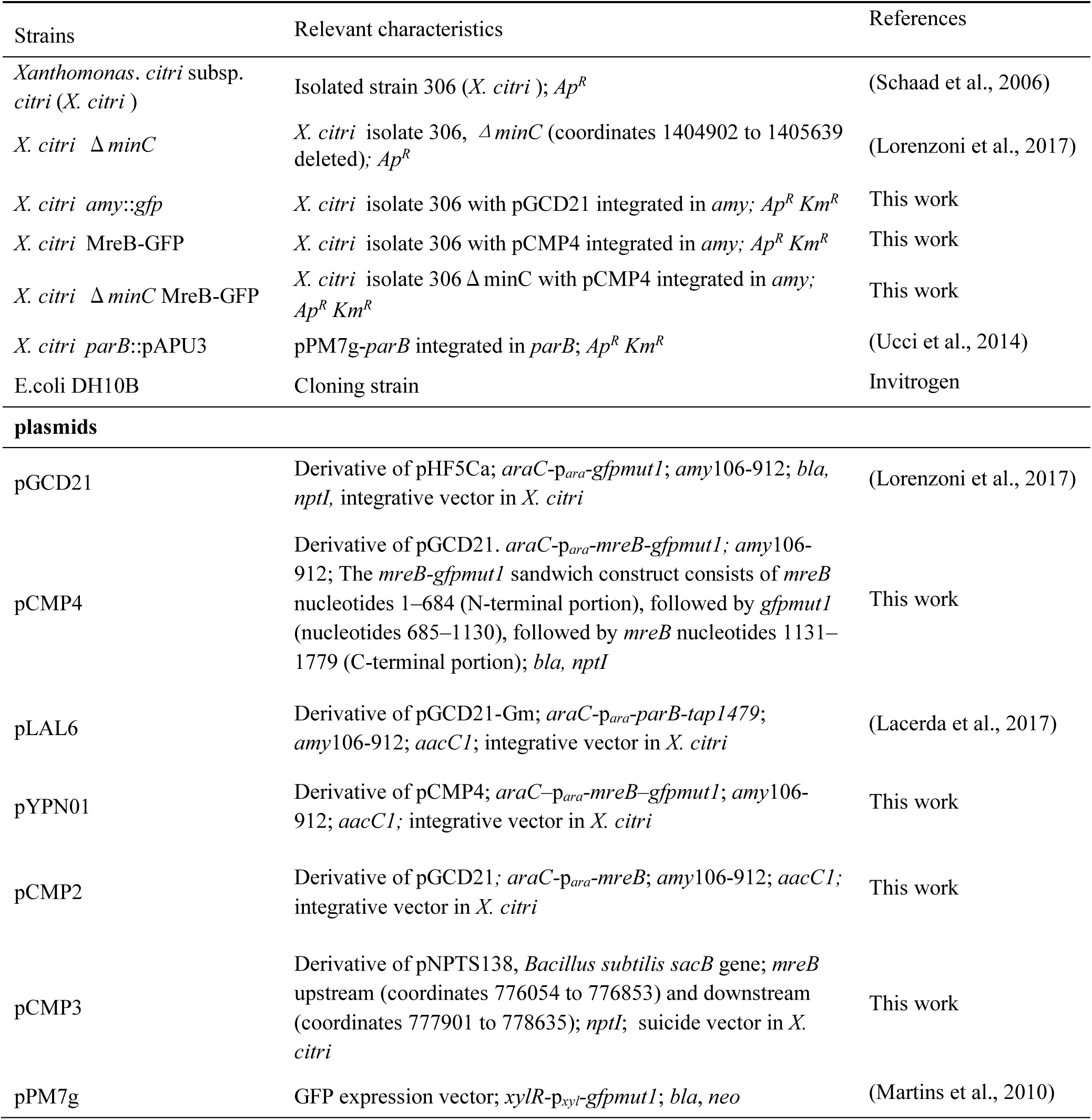
List of strains and plasmids.

### 2.2 Transformation

Both *X. citri* MreB-GFP and *X. citri* ΔminC MreB-GFP were generated by transformation of pCMP4 into the respective parent strains by electroporation (Ferreira et al., 1995).

### 2.3 Microscopy

Bacterial cells (1 μL) from the exponential growth phase were mounted on agarose pads (0.9% agarose in PBS) and observed using a Zeiss Axio Observer inverted microscope with a 100×/1.40 NA oil immersion objective. Illumination was provided by a Colibri 7 LED light source, and fluorescence was detected using appropriate filter sets for eGFP, and At425. All imaging was performed at room temperature. HADA (Hydroxycoumarin-carboxylic acid-Amino-D Alanine) is a fluorescent D-amino acid analogue that is incorporated into growing PG synthesis process, thereby labeling sites of PG formation (Peters et al., 2019). HADA was synthesized in our lab (Lorenzoni et al., 2017) and added to exponential phase *X. citri* cells at a final concentration of 250 µM, and the labeling was performed for 1 hour. A22 hydrochloride (Sigma-Aldrich SML0471, ≥95% purity (HPLC)) was added to exponential phase *X. citri* cells to a final concentration of 20 µg/mL. To visualize chromosomal DNA in live cells, cells are labeled with 4,6-diamidino-2-phenylindole (DAPI) following the staining protocol of Howell et al (2017).equations should be inserted in editable format from the equation editor.

### 2.4 Analysis

Microscope images were processed by ImageJ, a MicrobeJ plugin, and JACoP (Bolte & Cordelières, 2006; Ducret et al., 2016; Schindelin et al., 2012). Other data were analysed by using GraphPad Prism version 10.4.0 for Windows, GraphPad Software, Boston, Massachusetts USA, www.graphpad.com.

## 3 Results

### 3.1 The *X. citri* Δ *minC* MreB-GFP mutants display the reported Δ *minC* phenotype, which is growth medium dependent

In order to link cell wall synthesis to MreB localization, a *mreB-gfp* sandwich fusion was introduced as an extra copy at the *amy* locus of isolate 306 and also into the *X. citri* Δ*minC* mutant. As previously reported (Lorenzoni et al., 2017), deletion of *minC* in *X. citri* results in distinct phenotypic abnormalities, including branching cells, minicells, and elongated filaments (Fig. 1A). This phenotype is growth medium-dependent, with a higher frequency of branching cells on Xam1 medium compared to the nutrient-rich NYGB medium (Table 2). The nutrient limitation in Xam1 imposes growth stress, leading to impaired cell division. Introduction of MreB-GFP did not alter the length of cells compared to those of the wild type parental strain (Fig. 1B). Bacteria grown on NYGB medium werelonger (*p*< 0.001) compared to those grown on Xam1 medium (Fig. 1B), consistent with the observation that cell-length is correlated with nutrient availability in the medium (Schaechter et al., 1958).

**Fig. 1.**
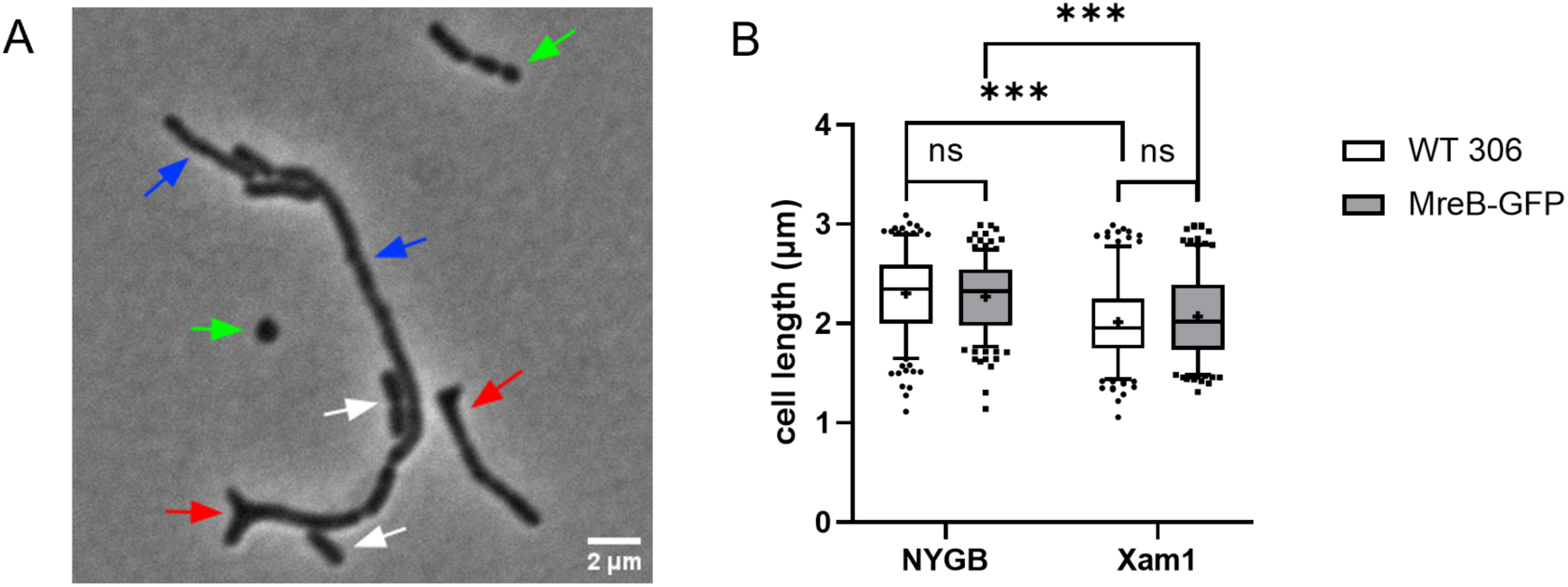
(A) Morphology of *X. citri ΔminC mreB-gfp* grown in Xam1 medium (White arrow: normal cells, red arrow: branching, green arrow: minicells, blue arrow: filaments). (B) Distribution of the cell lengths of wild type *X. citri* isolate 306 (WT 306) and *X. citri* isolate 306 expressing *mreB-gfp* (MreB-GFP) in NYGB and Xam1 medium (each group: 256 cells. Statistical significance, assumed when *p*-value <0.001, is marked with three asterisk (***), representing a cell length distribution that is significantly different from the other group).

**Table 2.** *X. citri* ΔminC MreB-GFP phenotypes in different media.

| Strains | Medium | Minicells | Branching | Short filaments | Normal | Total |
|---|---|---|---|---|---|---|
| *X. citri* ΔminC MreB-GFP | NYGB | 11.28% | 1.55% | 7.74% | 79.65% | 452 |
|  | Xam1 | 7.92% | 12.32% | 18.31% | 61.33% | 568 |
| *X. citri* ΔminC | NYGB | 8.08% | 1.09% | 10.92% | 79.91% | 458 |
|  | Xam1 | 8.44% | 12.67% | 19.11% | 59.78% | 450 |

### 3.2 MreB-GFP localizes at the inner membrane of cells, while PG synthesis is mostly visible at the septum

We studied the spatial relationship between the cytoskeletal protein MreB and cell wall synthesis by observing the localization of MreB-GFP within cells, while using the fluorescent D-amino acid analogue HADA to label the active synthesis sites of PG in the cell wall. HADA is incorporated into PG in the periplasm through the activity of D,D-transpeptidases or L,D-transpeptidases, rather than through cytoplasmic incorporation into the precursor molecule LipidII (Kuru et al., 2019).

MreB localizes at the inner membrane of living cells in both the wild type and in the Δ*minC* background (Fig. 2), consistent with MreB localization patterns described in other bacteria (Carballido-López et al., 2006; Garenne et al., 2020; Shiomi, 2017). This indicates that *minC* is not required for membrane localization of MreB. The signal occasionally appears more concentrated in one or two spots in the wild type background, compared to the *minC* deletion strain, but this may also have to do with the difference in cell appearance when cells start to branch and filament without *minC*. According to the geometric localization model, MreB is usually depleted at the poles and enriched in regions of low or negative mean curvature, which is the cylindrical region of the cell (Bratton et al., 2018). In contrast to the co-localization observed between cell division sites and PG synthesis activity (Lorenzoni et al., 2017), MreB exhibits a distinct spatial distribution pattern, that does not show overlap with HADA signal. As a dynamic cytoskeletal protein, MreB constantly moves within the cell, whereas HADA, once incorporated into the cell wall during PG synthesis, remains fixed at those sites. The HADA signal is mostly localized at the septum, indicative of active synthesis of PG at this site during cell division. However, it should be noted that this is also the site where the signal would be most concentrated and therefore most easily visible – some disperse HADA signal, indicative of PG synthesis along the lateral wall, can also be detected. Additionally, a small number of signals appear at the cell poles, likely because these regions have recently undergone division and represent newly formed poles. Importantly, after deletion of *minC*, transverse bands of HADA are still visible indicative of possible division sites. Earlier we established that these sites do no longer overlap with the localization of cell division proteins (Lorenzoni et al., 2017). It is quite evident that MreB does not accumulate at these sites either (Fig. 2B).

**Fig. 2.**
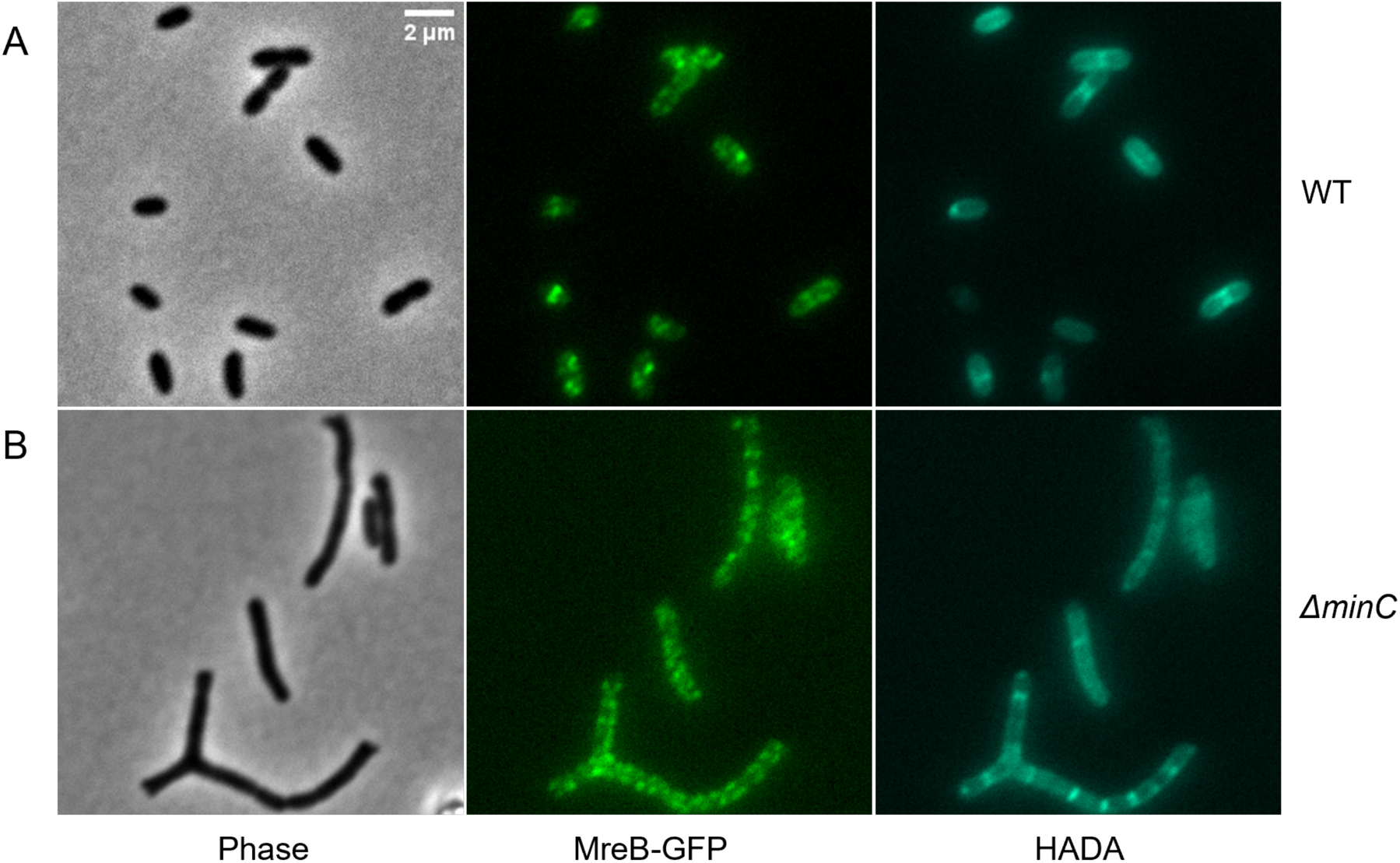
Localization of MreB-GFP and peptidoglycan synthesis in *X. citri mreB-gfp* (A) and *X. citri ΔminC mreB-gfp* (B) in Xam1. Phase contrast (left), GFP (center), and HADA (right, peptidoglycan incorporation sites) images were taken with the appropriate filter sets.

We noticed that MreB-GFP localized in bright polar spots in cells grown at 30°C (Fig. S1A). These spots are caused by formation of inclusion bodies containing MreB-GFP, as evidenced by a mild detergent extraction experiment (Fig. S1B). The bright polar spots disappeared, and inclusion body formation was undetectable when cells were grown at 25 °C (Fig. S1), therefore all localization experiments including the one described above were performed at 25 °C.

### 3.3 Treatment with the MreB inhibitor A22 leads to loss of rod shape and inhibition of cell growth

To determine whether PG synthesis is dependent on MreB, we employed the MreB-perturbing compound A22 (S-(3,4-dichlorobenzyl) isothiourea). A22 is an inhibitor of MreB that disrupts MreB polymer assembly by competitively binding to MreB monomers and blocking ATP binding (Bonez et al., 2016; Wang et al., 2010). A22 induced a gradual morphological change. After exposure to 20 μg/mL A22 for 4 hours, cells exhibited progressive shortening and widening, ultimately losing their rod-shaped characteristics after around 15 hours of continued growth (Fig. 3A, B). Following A22 treatment, the spherical cells of the Δ*minC* MreB-GFP strain were considerably larger than those of *X. citri* MreB-GFP, consistent with the observation that the Δ*minC* mutant displayed a greater average cell length before treatment. Growth curves of cultures clearly showed that the inhibitory effect of A22 on *X. citri* cell growth and division is dose-dependent (Fig. S2), and that the Δ*minC* mutant is much more sensitive to A22 (Fig. 3C). This strongly suggests that MreB-linked PG synthesis becomes critical when cell division is perturbed in the absence of MinC.

**Fig. 3.**
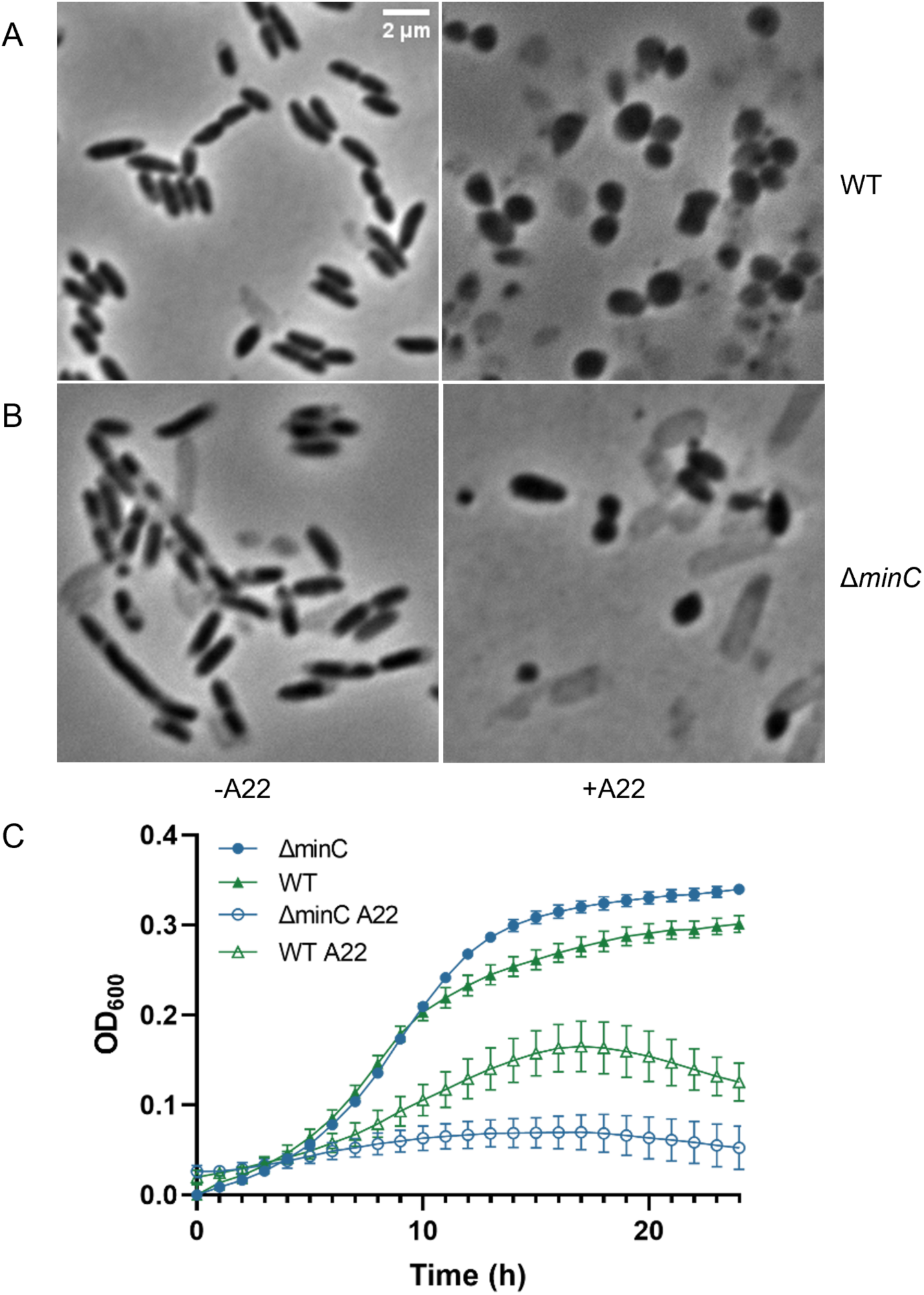
A22 treatment induces rounding of cells. Cells of *X. citri* MreB-GFP (A) and *X. citri* Δ*minC* MreB-GFP (B) were grown in NYGB for 15 h in the absence (left) or presence of 20μg/mL A22 (right). (C) Growth curves of *X. citri* cells grown with or without A22 (20 μg/mL) in NYGB.

### 3.4 A22 treatment induces the reversible delocalization of MreB to the cell cap regions, where it is partially co-localized with PG synthesis sites

The rod cell consists of two main sections: a central cylindrical region and rounded caps located at each end of the cylinder (Shiomi et al., 2009). Many studies have demonstrated that loss of MreB function is lethal and pleiotropic, affecting various essential cellular processes including cell shape maintenance, cell division, and chromosome segregation (Gitai et al., 2005; Shi et al., 2018). We observed that non-functional MreB accumulated as small fluorescent clusters specifically at the cell cap regions (Fig. 4A, B), in agreement with the *in vitro* MreB behavior under A22 treatment reported by Bean et al (2009). The polar localization pattern is readily reversible by removal of A22 which resulted in MreB redistribution within 30 min after removal (Fig. 4C). A demographic analysis of MreB-GFP signal distribution, shows that MreB-GFP shifts from a uniform or central distribution to the cell polar region upon A22 treatment, and reverses back after A22 washout (Fig. 4D). The reversibility of the A22 effect suggested that the fluorescent polar signal was not due to the formation of insoluble inclusion bodies, which was confirmed by a detergent extraction experiment in the presence of A22 (Fig. S1).

**Fig. 4.**
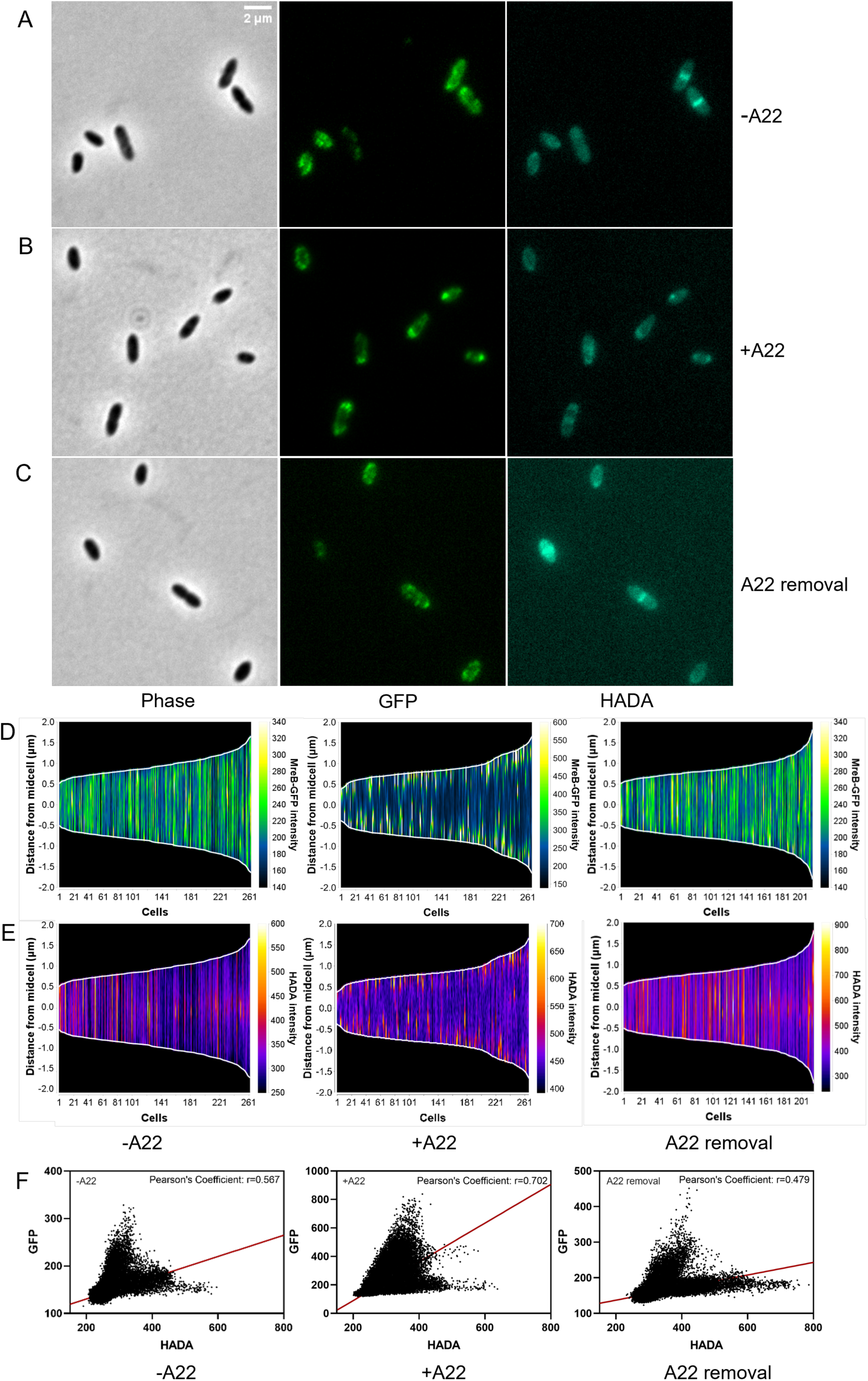
A22 treatment causes MreB-GFP to localize in polar clusters. Cells were either not treated with A22 (control, panel A), treated with A22 (20 µg/mL) for 4 hours (panel B), or treated with A22 (20 µg/mL) for 4 hours followed by removal of A22 by washing and growth for an additional 30 mins (panel C). Phase contrast (left). GFP (center) and HADA labelling (right) were imaged. (D-E) Demographs of MreB-GFP (D) and HADA (E) signal distribution, generated with MicrobeJ in ImageJ and aligned by cell length. (F) Co-localization analysis of MreB-GFP and HADA signals, quantified using Pearson’s correlation (JACoP in ImageJ). Panels show representative data for untreated (left, n=262 cells), A22-treated (20 µg/mL; center, n=266 cells), and A22 washout (right, n=214 cells) conditions.

After A22 treatment, PG synthesis sites begin to partially overlap with MreB, with some of the newly generated PG no longer located at division septa but instead appearing at the cell poles (Fig. 4E). The high incidence and reversibility of polar PG synthesis following A22 removal, along with the Pearson’s correlation analysis of the HADA and GFP signals (Fig. 4F), argue that the appearance of polar HADA spots is caused by active, MreB-linked, PG synthesis. This suggests that MreB delocalization disrupts the spatial regulation of PG insertion, leading to synthesis at atypical polar sites.

### 3.5 MreB is required for proper ParABS-mediated chromosome segregation

To assess the impact of MreB dysfunction on chromosome segregation, we studied the distribution of ParB-GFP, which binds to specific *parS* sites on the chromosome, in A22 treated cells (Ucci et al., 2014). ParB localization can be used as a measure for equitable distribution of genetic material during cell division (Chu et al., 2024; Jalal and Le, 2020). In growing cells, ParB-GFP could be found to be localizing to the cell poles in longer cells (Fig. 5A), indicating correct segregation of the replicated chromosomes prior to division, as previously described (Ucci et al., 2014). After 2 h of A22 treatment, the ParB-GFP signal started to appear dispersed, and a low number of anucleate cells could be detected (Fig 5B, Table 3). This result indicates that correct MreB functioning is important for chromosome segregation.

**Fig. 5.**
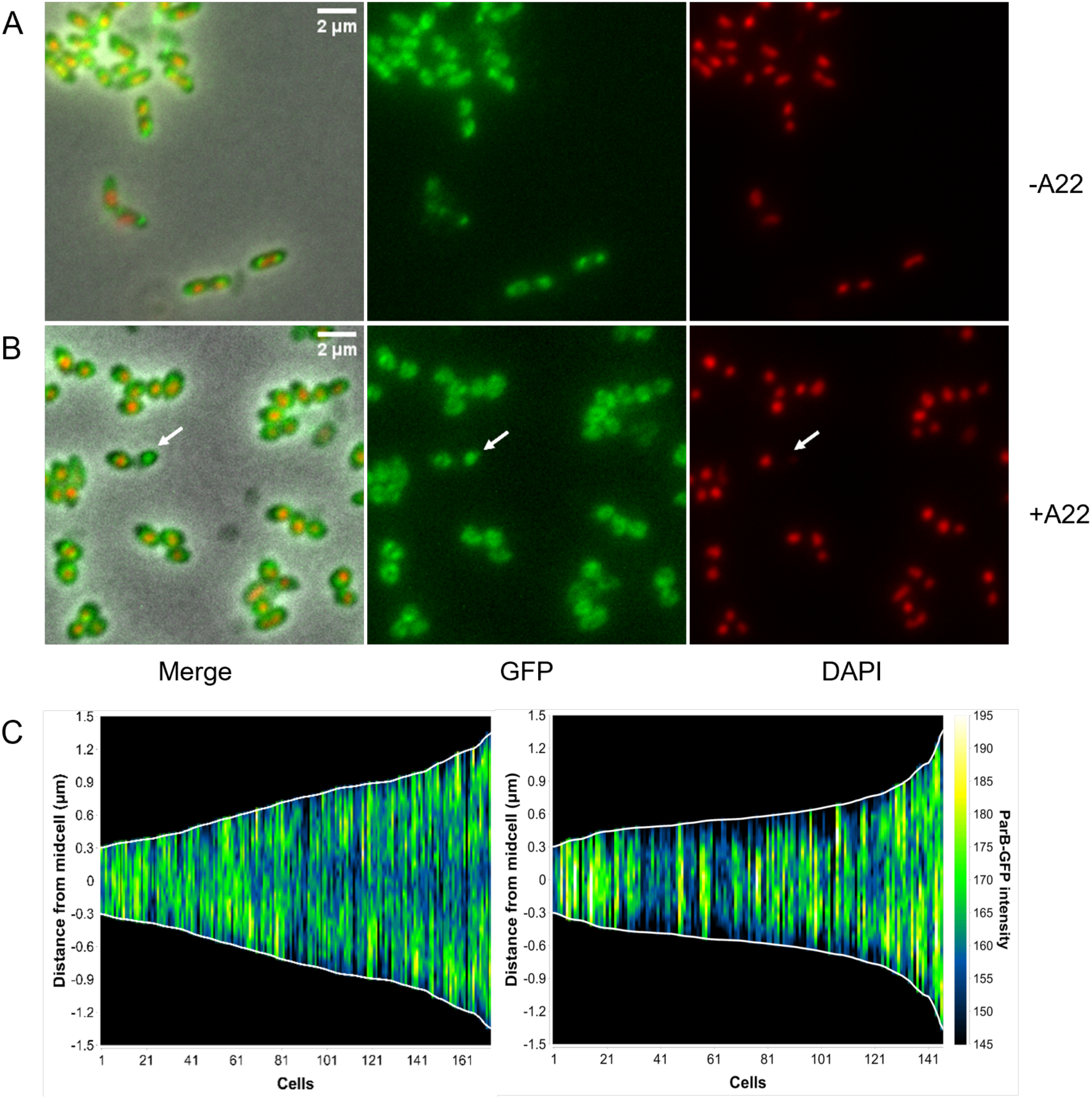
ParB distribution is affected by A22 treatment. *X. citri parB-gfp* cells were grown for 2 h, without (A) or with 20 µg/mL A22 (B), stained with DAPI and phase contrast, GFP and DAPI images were taken. (C) Heat maps of ParB-GFP distribution were organized based on cell length ((left): without A22 treatment, n=173, (right): 20 μg/mL A22 treatment, n=146) and stacked to create demographs using MicrobeJ in ImageJ.

**Table 3.**
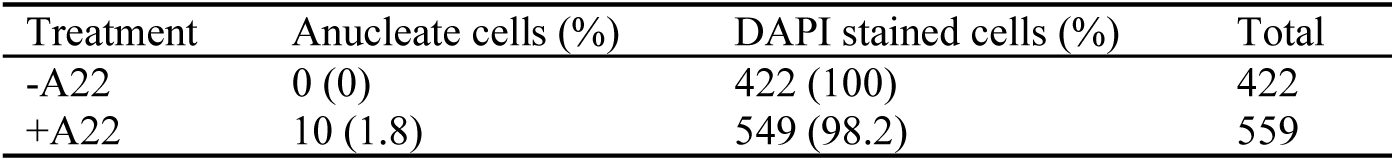
Anucleate cells percentage in *X. citri* ParB-GFP as determined using DAPI staining.

## 4 Discussion

In this study, we investigated the role of MreB in shape determination and chromosome segregation in *Xanthomonas citri*. We previously showed that during aberrant septum positioning in a *X. citri minC* deletion strain, the localization of cell division proteins appeared uncoupled from PG synthesis (Lorenzoni et al., 2017). In this paper, we explored whether MreB localizes to these ‘novel’ PG synthesis sites in a Δ*minC* mutant. We found that MreB is not accumulating at aberrantly placed septa (Fig. 2). Nevertheless, MreB dependent PG synthesis becomes more critical when septum placement is affected as can be seen from the increased sensitivity to A22 in the Δ*minC* mutant (Fig. 3). This indicates that MreB driven PG synthesis becomes critical in the absence of divisome regulation, but exactly how is still unclear. A caveat is that the localization of division proteins over the course of division is rather static – always at the septum, which may narrow but stays put – whereas MreB is more mobile in the cell, moving along the entire lateral wall. Since HADA labelling does not provide a snapshot of PG synthesis but rather reports the aggregate of the places where HADA has been incorporated over the duration of the labeling period, colocalization of proteins to a HADA stained septum is easier to detect than colocalizing HADA with a moving elongasome.

The latter is also evident from the A22 experiment – A22 caused both MreB and PG synthesis to delocalize to cell poles (Fig. 4). This more static localization of MreB may have allowed sufficient HADA labelling at the same site to show colocalization. Prolonged aberrant localization of MreB and PG synthesis in the presence of A22 resulted in loss of shape and, ultimately, lysis.

To assess the physiological necessity of MreB, and the functionality of our MreB-GFP fusion, we attempted to generate a chromosomal *mreB* deletion mutant (see supplemental material) using a *sacB* suicide vector. We were unable to generate such a mutant in either a wild type background or in strains that conditionally express *mreB* or *mreB-gfp* from *amy*. Sequencing of false positives indicated retention of *mreB*. Although technical factors, including *sacB* counter-selection instability, or the necessity of the entire *mreBCD* operon for complementation (Jones et al., 2001), rather than *mreB* alone, cannot be ruled out as the cause of the failed knockout, the consistent failure across multiple independent attempts, together with the dose-dependent growth inhibition by A22, strongly indicate that *mreB* is an essential gene in *X. citri*.

A22 treatment triggered a distinct and reversible redistribution of MreB to the cell poles (Fig. 4). Sun et al. (2025) demonstrated that a high-osmolarity shock induced by 1M sorbitol drives the rapid and reversible relocalization of MreB to the cell poles and polar cell wall synthesis in *E. coli*. This would provide new physical linkages at the pole, between the cytoplasmic membranes and the cell wall, thereby limiting cytoplasmic contraction under osmotic stress. While the underlying mechanisms are likely distinct, A22 disrupts MreB polymerization through competitive ATP binding, whereas osmotic stress imposes physical stress on the linkage between peptidoglycan and the inner membrane, the similarity raises the possibility that polar recruitment of MreB may represent a conserved cellular response to stress in the peptidoglycan-membrane connection, regardless of its origin. The structural integrity and spatial organization of MreB filaments are sensitive to environmental fluctuations. In *B. subtilis*, increased osmotic pressure triggers MreB-GFP to transition from a dispersed filaments and spots into a less but brighter structures, with a concomitant shift in PG synthesis (Dersch et al., 2020). Beyond osmotic and chemical stress, MreB filaments exhibit significant sensitivity to physiological stressors: non-aerated incubation leads to the dissolution of filaments into small stationary foci, while heat shock (50°C) induces the formation of shorter and disordered filament structures (Dersch et al., 2020). Altogether, these data highlight MreB cytoskeleton as a highly responsive sensor that undergoes rapid structural adaptation to diverse environmental challenges.

Our findings also provide evidence that a functional MreB cytoskeleton is important for robust ParABS-mediated chromosome segregation in *X. citri* (Fig. 5). In *E. coli*, which lacks chromosomal *parABS*, inhibition of MreB by A22 caused mislocalization of plasmid encoded ParB-GFP, as well as the appearance of anucleate cells, a phenotype indicative of failed chromosome partitioning (Iwai et al., 2002; Kruse et al., 2003). A direct role for MreB in chromosome segregation in *E. coli* has been contested (Karczmarek et al., 2007). Soufo and Graumann (2003) reported that in MreB or Mbl-depleted *B. subtilis*, anucleate cells emerged, the orientation of the origin regions became disordered, and chromosomes migrated more randomly or even failed to segregate.

Several explanations for the need for MreB for correct chromosome segregation have been put forward. First, the MreB cytoskeleton may act as a spatial organizer or a track that actively guides the directional movement of the ParB-*parS* nucleoprotein complex and chromosomal DNA in opposite directions in co-ordination with lateral cell wall growth (Kruse et al., 2003). In this explanation, ParA-mediated ATP hydrolysis provides the driving force, but MreB structures ensure this force is applied along the long cellular axis to achieve separation toward the poles. The dissolution of this track upon A22 treatment would cause the ParB complexes to lose their directional guidance, leading to the observed dispersion and failed segregation. In organisms lacking ParABS, undiscovered motor-like proteins or coupling factors might utilize MreB filaments to facilitate poleward chromosome separation (Soufo and Graumann, 2003; van Teeffelen et al., 2011). A second possibility is that the loss of the rod shape of the cell perturbs entropy-driven chromosome segregation. In a cylindrical cell, entropic polymer-repulsion forces between duplicated chromosomes naturally drive their spontaneous segregation along the longitudinal axis (Jun and Mulder, 2006; Jun and Wright, 2010). Consequently, cell rounding caused by A22-mediated MreB disruption would interfere with this geometric confinement, and ultimately culminates in defective chromosome segregation. Distinguishing between these models will require future experiments such as single-molecule tracking of ParB under MreB perturbation, or identification of potential MreB-associated motor-like proteins in *X. citri*.

Interestingly, we observed that anucleate cells lacked detectable DAPI signal yet retained ParB-GFP fluorescence (Fig. 5). This could be due to the accumulation of free ParB following overexpression. Cellular ParB exists either in a *parS*-bound pool or a freely diffusing cytoplasmic pool. Lim et al. demonstrated that GFP-ParB/*parS* foci account for ∼80% of total GFP-ParB signal, whereas the remaining ∼20% corresponds to diffusing protein and potential degradation products (Lim et al., 2014). The disruption of chromosome separation coordination may also cause guillotining of the chromosome, with a DNA fragment that is too small to be detected using DAPI, containing (Iwai et al., 2002)*parS*/ParB ending up in a daughter cell.

In summary, this study provides a characterization of MreB function in *Xanthomonas*. We demonstrate that MreB maintains its inner membrane localization independently of MinC, while its dysfunction through A22 treatment leads to progressive morphological defects, mislocalized PG synthesis, and impaired chromosome segregation. The failure to generate a viable *mreB* deletion mutant, together with the dose-dependent growth inhibition by A22, strongly supports the essentiality of MreB in *X. citri*. Collectively, our findings highlight MreB as a central coordinator of multiple cellular processes, such as cell shape maintenance, cell wall synthesis, and chromosome segregation, as it is in other bacteria.

## 6 Author Contributions

YP: Data curation, Formal analysis, Funding acquisition, Investigation, Methodology, Software, Validation, Visualization, Writing – original draft, Writing – review & editing. TV: Data curation, Formal analysis, Investigation, Validation, Visualization, Writing – original draft, Writing – review & editing. CMP: Data curation, Funding acquisition, Investigation, Methodology, Validation. HF: Conceptualization, Formal analysis, Funding acquisition, Resources, Methodology, Validation, Writing – review & editing. DJS: Conceptualization, Data curation, Formal analysis, Funding acquisition, Investigation, Methodology, Validation, Writing – original draft, Writing – review & editing.

## 8 Data Availability Statement

The original contributions presented in the study are publicly available. The whole genome sequencing data obtained in this study have been submitted to SRA with the accession number PRJNA1522837.

## References

1. Ali, S., Hameed, A., Muhae-Ud-Din, G., Ikhlaq, M., Ashfaq, M., Atiq, M., Ali, F., Zia, Z. U., Naqvi, S. A. H., Wang, Y. (2023). Citrus Canker: A Persistent Threat to the Worldwide Citrus Industry—An Analysis. Agronomy, 13(4), 1112. https://www.mdpi.com/2073-4395/13/4/1112

2. Barrows, J. M., Goley, E. D. (2021). FtsZ dynamics in bacterial division: What, how, and why? Current Opinion in Cell Biology, 68, 163–172. 10.1016/j.ceb.2020.10.013

3. Bean, G. J., Flickinger, S. T., Westler, W. M., McCully, M. E., Sept, D., Weibel, D. B., Amann, K. J. (2009). A22 disrupts the bacterial actin cytoskeleton by directly binding and inducing a low-affinity state in MreB. Biochemistry, 48(22), 4852–4857. 10.1021/bi900014d

4. Bock, F. P., Liu, H. W., Anchimiuk, A., Diebold-Durand, M.-L., Gruber, S. (2022). A joint-ParB interface promotes Smc DNA recruitment. Cell Reports, 40(9). 10.1016/j.celrep.2022.111273

5. Böhm, K., Giacomelli, G., Schmidt, A., Imhof, A., Koszul, R., Marbouty, M., Bramkamp, M. (2020). Chromosome organization by a conserved condensin-ParB system in the actinobacterium Corynebacterium glutamicum. Nature Communications, 11(1), 1485. 10.1038/s41467-020-15238-4

6. Bonez, P. C., Ramos, A. P., Nascimento, K., Copetti, P. M., Souza, M. E., Rossi, G. G., Agertt, V. A., Sagrillo, M. R., Santos, R. C., Campos, M. M. (2016). Antibacterial, cyto and genotoxic activities of A22 compound ((S-3, 4-dichlorobenzyl) isothiourea hydrochloride). Microb Pathog, 99, 14–18. 10.1016/j.micpath.2016.07.007

7. Bottomley, A. L., Liew, A. T. F., Kusuma, K. D., Peterson, E., Seidel, L., Foster, S. J., Harry, E. J. (2017). Coordination of Chromosome Segregation and Cell Division in Staphylococcus aureus. Front Microbiol, 8, 1575. 10.3389/fmicb.2017.01575

8. Bratton, B. P., Shaevitz, J. W., Gitai, Z., Morgenstein, R. M. (2018). MreB polymers and curvature localization are enhanced by RodZ and predict E. coli’s cylindrical uniformity. Nature Communications, 9(1), 2797. 10.1038/s41467-018-05186-5

9. Cameron, T. A., Margolin, W. (2024). Insights into the assembly and regulation of the bacterial divisome. Nature Reviews Microbiology, 22(1), 33–45. 10.1038/s41579-023-00942-x

10. Cao, X. Q., Ouyang, X. Y., Chen, B., Song, K., Zhou, L., Jiang, B. L., Tang, J. L., Ji, G., Poplawsky, A. R., He, Y. W. (2020). Genetic Interference Analysis Reveals that Both 3-Hydroxybenzoic Acid and 4-Hydroxybenzoic Acid Are Involved in Xanthomonadin Biosynthesis in the Phytopathogen Xanthomonas campestris pv. campestris. Phytopathology®, 110(2), 278–286. 10.1094/phyto-08-19-0299-r

11. Carballido-López, R., Formstone, A., Li, Y., Ehrlich, S. D., Noirot, P., Errington, J. (2006). Actin Homolog MreBH Governs Cell Morphogenesis by Localization of the Cell Wall Hydrolase LytE. Developmental Cell, 11(3), 399–409. 10.1016/j.devcel.2006.07.017

12. Chu, C. H., Wu, C. T., Lin, M. G., Yen, C. Y., Wu, Y. Z., Hsiao, C. D., Sun, Y. J. (2024). Insights into the molecular mechanism of ParABS system in chromosome partition by HpParA and HpParB. Nucleic Acids Research, 52(12), 7321–7336. 10.1093/nar/gkae450

13. Dersch, S., Mehl, J., Stuckenschneider, L., Mayer, B., Roth, J., Rohrbach, A., Graumann, P. L. (2020). Super-Resolution Microscopy and Single-Molecule Tracking Reveal Distinct Adaptive Dynamics of MreB and of Cell Wall-Synthesis Enzymes. Frontiers in Microbiology, Volume 11 - 2020. 10.3389/fmicb.2020.01946

14. Domínguez-Escobar, J., Chastanet, A., Crevenna, A. H., Fromion, V., Wedlich-Söldner, R., Carballido-López, R. (2011). Processive Movement of MreB-Associated Cell Wall Biosynthetic Complexes in Bacteria. Science, 333(6039), 225–228. doi:10.1126/science.1203466

15. Ference, C. M., Gochez, A. M., Behlau, F., Wang, N., Graham, J. H., Jones, J. B. (2018). Recent advances in the understanding of Xanthomonas citri ssp. citri pathogenesis and citrus canker disease management. Molecular Plant Pathology, 19(6), 1302–1318. 10.1111/mpp.12638

16. Ferreira, H., Barrientos, F. J. A., Baldini, R. L., Rosato, Y. B. (1995). Electrotransformation of three pathovars of Xanthomonas campestris. Applied Microbiology and Biotechnology, 43(4), 651–655. 10.1007/BF00164769

17. Figge, R. M., Divakaruni, A. V., Gober, J. W. (2004). MreB, the cell shape-determining bacterial actin homologue, co-ordinates cell wall morphogenesis in Caulobacter crescentus. Molecular Microbiology, 51(5), 1321–1332. 10.1111/j.1365-2958.2003.03936.x

18. Funnell, B. E. (2016). ParB Partition Proteins: Complex Formation and Spreading at Bacterial and Plasmid Centromeres [Mini Review]. Frontiers in Molecular Biosciences, Volume 3 - 2016. 10.3389/fmolb.2016.00044

19. Garenne, D., Libchaber, A., Noireaux, V. (2020). Membrane molecular crowding enhances MreB polymerization to shape synthetic cells from spheres to rods. Proceedings of the National Academy of Sciences, 117(4), 1902–1909. doi:10.1073/pnas.1914656117

20. Ghosal, D., Trambaiolo, D., Amos, L. A., Löwe, J. (2014). MinCD cell division proteins form alternating copolymeric cytomotive filaments. Nature Communications, 5(1), 5341. 10.1038/ncomms6341

21. Gitai, Z., Dye, N. A., Reisenauer, A., Wachi, M., Shapiro, L. (2005). MreB Actin-Mediated Segregation of a Specific Region of a Bacterial Chromosome. Cell, 120(3), 329–341. 10.1016/j.cell.2005.01.007

22. Grinnell, A., Sloan, R., Morgenstein, R. M. (2022). Cell density-dependent antibiotic tolerance to inhibition of the elongation machinery requires fully functional PBP1B. Commun Biol, 5(1), 107. 10.1038/s42003-022-03056-x

23. Iwai, N., Nagai, K., Wachi, M. (2002). Novel S-benzylisothiourea compound that induces spherical cells in Escherichia coli probably by acting on a rod-shape-determining protein(s) other than penicillin-binding protein 2. Biosci Biotechnol Biochem, 66(12), 2658–2662. 10.1271/bbb.66.2658

24. Jalal, A. S. B., Le, T. B. K. (2020). Bacterial chromosome segregation by the ParABS system. Open Biology, 10(6), 200097. doi:10.1098/rsob.200097

25. Jones, L. J. F., Carballido-López, R., Errington, J. (2001). Control of Cell Shape in Bacteria. Cell, 104(6), 913–922. 10.1016/S0092-8674(01)00287-2

26. Jun, S., Mulder, B. (2006). Entropy-driven spatial organization of highly confined polymers: Lessons for the bacterial chromosome. Proceedings of the National Academy of Sciences, 103(33), 12388–12393. doi:10.1073/pnas.0605305103

27. Jun, S., Wright, A. (2010). Entropy as the driver of chromosome segregation. Nature Reviews Microbiology, 8(8), 600–607. 10.1038/nrmicro2391

28. Karczmarek, A., Martínez-Arteaga, R., Alexeeva, S., Hansen, F. G., Vicente, M., Nanninga, N., den Blaauwen, T. (2007). DNA and origin region segregation are not affected by the transition from rod to sphere after inhibition of Escherichia coli MreB by A22. Mol Microbiol, 65(1), 51–63. 10.1111/j.1365-2958.2007.05777.x

29. Kruse, T., Møller-Jensen, J., Løbner-Olesen, A., Gerdes, K. (2003). Dysfunctional MreB inhibits chromosome segregation in *Escherichia coli*. The EMBO Journal, 22(19), 5283–5292. 10.1093/emboj/cdg504

30. Kuru, E., Radkov, A., Meng, X., Egan, A., Alvarez, L., Dowson, A., Booher, G., Breukink, E., Roper, D. I., Cava, F., Vollmer, W., Brun, Y., VanNieuwenhze, M. S. (2019). Mechanisms of Incorporation for D-Amino Acid Probes That Target Peptidoglycan Biosynthesis. ACS Chemical Biology, 14(12), 2745–2756. 10.1021/acschembio.9b00664

31. Lacerda, L. A., Cavalca, L. B., Martins, P. M. M., Govone, J. S., Bacci, M., Ferreira, H. (2017). Protein depletion using the arabinose promoter in Xanthomonas citri subsp. citri. Plasmid, 90, 44–52. 10.1016/j.plasmid.2017.03.005

32. Lanza, F. E., Marti, W., Silva, G. J., Behlau, F. (2019). Characteristics of Citrus Canker Lesions Associated with Premature Drop of Sweet Orange Fruit. Phytopathology, 109(1), 44–51. 10.1094/phyto-04-18-0114-r

33. Lim, H. C., Surovtsev, I. V., Beltran, B. G., Huang, F., Bewersdorf, J., Jacobs-Wagner, C. (2014). Evidence for a DNA-relay mechanism in ParABS-mediated chromosome segregation. eLife, 3, e02758. 10.7554/eLife.02758

34. Liu, L., Liu, X., Liu, L., Zhu, T., Ye, R., Chen, H., Zhou, L., Wu, G., Tan, L., Han, J., Li, R., Ma, X., Deng, Z. (2024). Clarification of the infection pattern of Xanthomonas citri subsp. citri on citrus fruit by artificial inoculation. Plant Methods, 20(1), 65. 10.1186/s13007-024-01190-7

35. Livny, J., Yamaichi, Y., Waldor Matthew, K. (2007). Distribution of Centromere-Like parS Sites in Bacteria: Insights from Comparative Genomics. Journal of bacteriology, 189(23), 8693–8703. 10.1128/jb.01239-07

36. Lorenzoni, A. S. G., Dantas, G. C., Bergsma, T., Ferreira, H., Scheffers, D.-J. (2017). Xanthomonas citri MinC Oscillates from Pole to Pole to Ensure Proper Cell Division and Shape. Frontiers in Microbiology, 8, 1352. 10.3389/fmicb.2017.01352

37. Martins, P. M., Lau, I. F., Bacci, M., Belasque, J., do Amaral, A. M., Taboga, S. R., Ferreira, H. (2010). Subcellular localization of proteins labeled with GFP in Xanthomonas citri ssp. citri: targeting the division septum. FEMS Microbiol Lett, 310(1), 76–83. 10.1111/j.1574-6968.2010.02047.x

38. Parham, J., Sorichetti, V., Cezanne, A., Foo, S., Kuo, Y. W., Hoogenberg, B., Radoux-Mergault, A., Mawdesley, E., Gatward, L. D., Boulanger, J., Schulze, U., Šarić, A., Baum, B. (2025). Temporal and spatial coordination of DNA segregation and cell division in an archaeon. Proc Natl Acad Sci U S A, 122(42), e2513939122. 10.1073/pnas.2513939122

39. Rajagopal, L., Sundari, C. S., Balasubramanian, D., Sonti, R. V. (1997). The bacterial pigment xanthomonadin offers protection against photodamage. FEBS Letters, 415(2), 125–128. 10.1016/S0014-5793(97)01109-5

40. Rothfield, L., Taghbalout, A., Shih, Y.-L. (2005). Spatial control of bacterial division-site placement. Nature Reviews Microbiology, 3(12), 959–968. 10.1038/nrmicro1290

41. Schaad, N. W., Postnikova, E., Lacy, G., Sechler, A., Agarkova, I., Stromberg, P. E., Stromberg, V. K., Vidaver, A. K. (2006). Emended classification of xanthomonad pathogens on citrus. Systematic and Applied Microbiology, 29(8), 690–695. 10.1016/j.syapm.2006.08.001

42. Schaechter, M., MaalØe, O., Kjeldgaard, N. O. (1958). Dependency on Medium and Temperature of Cell Size and Chemical Composition during Balanced Growth of Salmonella typhimurium. Microbiology, 19(3), 592–606. 10.1099/00221287-19-3-592

43. Shi, H., Bratton, B. P., Gitai, Z., Huang, K. C. (2018). How to Build a Bacterial Cell: MreB as the Foreman of E. coli Construction. Cell, 172(6), 1294–1305. 10.1016/j.cell.2018.02.050

44. Shiomi, D. (2017). Polar localization of MreB actin is inhibited by anionic phospholipids in the rod-shaped bacterium Escherichia coli. Current Genetics, 63(5), 845–848. 10.1007/s00294-017-0696-5

45. Shiomi, D., Mori, H., Niki, H. (2009). Genetic mechanism regulating bacterial cell shape and metabolism. Commun Integr Biol, 2(3), 219–220. 10.4161/cib.2.3.7930

46. Soufo, H. J. D., Graumann, P. L. (2003). Actin-like Proteins MreB and Mbl from *Bacillus subtilis* Are Required for Bipolar Positioning of Replication Origins. Current Biology, 13(21), 1916–1920. 10.1016/j.cub.2003.10.024

47. Sun, J., Cheng, M. M.-C., van Wijngaarden, E., Hernandez, C. J., Bernhardt, T. G., Huang, K. C. (2025). A load-bearing function for the cytoplasmic membrane of *Escherichia coli*. bioRxiv, 2025.2010.2002.680147. 10.1101/2025.10.02.680147

48. Szwedziak, P., Löwe, J. (2013). Do the divisome and elongasome share a common evolutionary past? Current Opinion in Microbiology, 16(6), 745–751. 10.1016/j.mib.2013.09.003

49. Takacs, C. N., Poggio, S., Charbon, G., Pucheault, M., Vollmer, W., Jacobs-Wagner, C. (2010). MreB Drives *De Novo* Rod Morphogenesis in *Caulobacter crescentus* via Remodeling of the Cell Wall. Journal of bacteriology, 192(6), 1671–1684. doi:10.1128/jb.01311-09

50. Tišma, M., Kaljević, J., Gruber, S., Le, T. B. K., Dekker, C. (2024). Connecting the dots: key insights on ParB for chromosome segregation from single-molecule studies. FEMS Microbiol Rev, 48(1). 10.1093/femsre/fuad067

51. Typas, A., Banzhaf, M., van den Berg van Saparoea, B., Verheul, J., Biboy, J., Nichols, R. J., Zietek, M., Beilharz, K., Kannenberg, K., von Rechenberg, M., Breukink, E., den Blaauwen, T., Gross, C. A., Vollmer, W. (2010). Regulation of peptidoglycan synthesis by outer-membrane proteins. Cell, 143(7), 1097–1109. 10.1016/j.cell.2010.11.038

52. Ucci, A. P., Martins, P. M. M., Lau, I. F., Bacci Jr, M., Belasque Jr, J., Ferreira, H. (2014). Asymmetric chromosome segregation in Xanthomonas citri ssp. citri. MicrobiologyOpen, 3(1), 29–41. 10.1002/mbo3.145

53. van Teeffelen, S., Wang, S., Furchtgott, L., Huang, K. C., Wingreen, N. S., Shaevitz, J. W., Gitai, Z. (2011). The bacterial actin MreB rotates, and rotation depends on cell-wall assembly. Proceedings of the National Academy of Sciences, 108(38), 15822–15827. doi:10.1073/pnas.1108999108

54. Vecchiarelli, A. G., Han, Y.-W., Tan, X., Mizuuchi, M., Ghirlando, R., Biertümpfel, C., Funnell, B. E., Mizuuchi, K. (2010). ATP control of dynamic P1 ParA–DNA interactions: a key role for the nucleoid in plasmid partition. Molecular Microbiology, 78(1), 78–91. 10.1111/j.1365-2958.2010.07314.x

55. Wang, S., Arellano-Santoyo, H., Combs, P. A., Shaevitz, J. W. (2010). Actin-like cytoskeleton filaments contribute to cell mechanics in bacteria. Proceedings of the National Academy of Sciences, 107(20), 9182–9185. doi:10.1073/pnas.0911517107

56. Wettmann, L., Kruse, K. (2018). The Min-protein oscillations in Escherichia coli: an example of self-organized cellular protein waves. Philos Trans R Soc Lond B Biol Sci, 373(1747). 10.1098/rstb.2017.0111

57. White, C. L., Gober, J. W. (2012). MreB: pilot or passenger of cell wall synthesis? Trends in Microbiology, 20(2), 74–79. 10.1016/j.tim.2011.11.004

