## supplemental materials for "Spatial coordination of peptidoglycan synthesis by MreB directs morphogenesis in *Xanthomonas citri*"

### Interplay between *minC*, MreB localization, and peptidoglycan synthesis in *Xanthomonas citri*

### 1 Supplementary text:

#### 1.1 Materials and Methods

##### 1.1.1 Inclusion bodies preparation

The bacterial cultures, imaged in supplemental figure 1, were used to test for the presence of inclusion bodies. 5 mL bacterial culture was harvested by centrifugation (5000 × *g*, 10 min, 4°C) and resuspended in resuspension buffer (20 mM Tris-HCl, pH 8.0, 0.2 M NaCl, 1 mM EDTA). Cell densities were normalized to identical OD600nm values across all samples to ensure comparable protein baselines. Cells were lysed via probe sonication on ice (20 s on / 20 s off, 5 cycles) and centrifugated (5000 × *g*, 10 min, 4°C). The resulting supernatant was collected as the soluble fraction. To extract the insoluble inclusion bodies, the pellet was resuspended in ice-cold isolation buffer containing mild detergent (20 mM Tris-HCl, pH 8.0, 0.5 M NaCl, 2% Tween-20, 2 M urea). To ensure the complete lysis of residual intact cells, samples were conducted to three rounds of sonication in total. The inclusion bodies were retained in the pellet after centrifugation. For Western blot, the initial soluble supernatant and the final detergent-resistant pellet (resuspended in an equivalent initial buffer volume) were mixed 1:1 with 2× SDS loading buffer (120 mM Tris-HCl, pH 6.8, 4% SDS, 20% glycerol, 200 mM DTT, 0.02% bromophenol blue), boiled at 95°C for 5 min, and either analyzed directly via SDS-PAGE / Western blotting or stored at -20°C. Western Blots were developed with polyclonal anti-GFP antibodies (SIGMA, G1544, 1:2500) as described by Lorenzoni et al (2017). Chemiluminescent signals were developed using the CDP-Star reagent (Roche).

##### 1.1.2 Δ*mreB* mutant construction

Allelic exchange, using the suicide plasmid pCMP3, was used to try to delete *mreB*. The deletion vector pCMP3 was constructed by cloning the upstream (coordinates 776054 to 776853)and downstream (coordinates 777901 to 778635) flanking regions of the *mreB* gene from Xac into pJET1.2 for sequencing. The upstream fragment (*Hind*III/*Kpn*I) and downstream fragment (*Eco*RI/*Kpn*I) were excised then purified, and ligated into the suicide vector pNPTS138 digested with *Hind*III and *Eco*RI. pCMP3 contains the homologous arms of MreB upstream and downstream, and was introduced into *X. citri* by electroporation. Single-crossover integrants were selected on kanamycin plates. To facilitate the second homologous recombination via pop-out, colonies were incubated in non-selection liquid medium, followed by counter-selection on 5% sucrose plates. Colonies were screened for *mreB* deletion by PCR, and successful deletion mutants were confirmed by Sanger sequencing (provided by Azenta Life Sciences).

#### 1.2 Results

##### 1.2.1 Temperature sensitivity of MreB-GFP folding in *X. citri*

This assay separates membrane-associated proteins and weak aggregates from inclusion bodies. Mild extraction with 2% Tween-20 and 2 M urea solubilizes membranes and peripheral proteins, but leaves dense inclusion bodies intact in the pellet. Inclusion bodies were formed at 30°C in *X. citri* MreB-GFP, (Fig. S1B, lane 1P), but this insoluble signal disappeared at 25°C (Fig. S1B, lane 4P). This is consistent with the microscopy results (Fig. S1A), confirming that the fluorescent foci observed under these conditions represent protein aggregates. The same assay was carried out in the presence of A22 to show that the GFP-bright spots formed after addition of A22 were not caused by the formation of inclusion bodies (Fig. S1B, lane 5P).

##### 1.2.2 Validation of *mreB* essentiality and knock-out attempts in *X. citri*

Colonies were selected using 3% sucrose plates as the colony PCR template. Wild type *X. citri* would show a band of 2452 bp (flanking region plus mreB gene), while MreB-deficient Xac should show approximately 1405 bp (lateral band only). Among 148 colonies of *X. citri* WT-pCMP3 screened, two were identified by diagnostic PCR as potential *mreB* deletion mutants. Both mutants retained kanamycin resistance, confirming the presence of the vector. Similarly, screening of 152 Xac pYPN01-pCMP3 colonies yielded two potential *mreB* deletion mutants. Whole-genome sequencing of the two candidate mutants revealed that the intended knockout was unsuccessful.(Fig. S3-4)

A hypothesis was proposed to explain results: there is the vector or part of it in the genome, which would allow the amplification of both the diagnostic band, along with some event that eliminated the lethal effect of *sacB* on *X. citri* in the presence of sucrose. Additionally, counter selection by *sacB* gene suffers from relatively high false positive rates for chromosomal manipulations due to spontaneous suppressor mutations that inactivate *sacB* (Khetrapal et al., 2015; Ma et al., 2015; Wang et al., 2019; Zhang et al., 1998).

The supplementation and deletion strategies with the suicide plasmid were earlier shown to be effective for the deletion of *parB* in *X. citri*, by Lacerda et al. (2017). However, *parB* is classified as a non-essential gene, which is an important distinction. We proposed the obtained results are most likely due to insufficient supplementation to ensure the vitality of the cells. In this case, the expression of *mreB* integrated into *amy* under the control of the promoter *arac* may be excessive or insufficient. It is also possible that certain features in the genomic region where *mreB* prevent or hinder deletion via double recombination. Jones et al. (2001) attempted to construct a merodiploid *Bacillus subtilis* strain with a knockout of *mreB* at its natural position and an inducible copy of *mreB* located at the *amyE* locus, but were unsuccessful. Only when the entire *mreBCD* cluster was positioned at *amyE*, *mreB* could be deleted.

#### 2 Supplementary figures

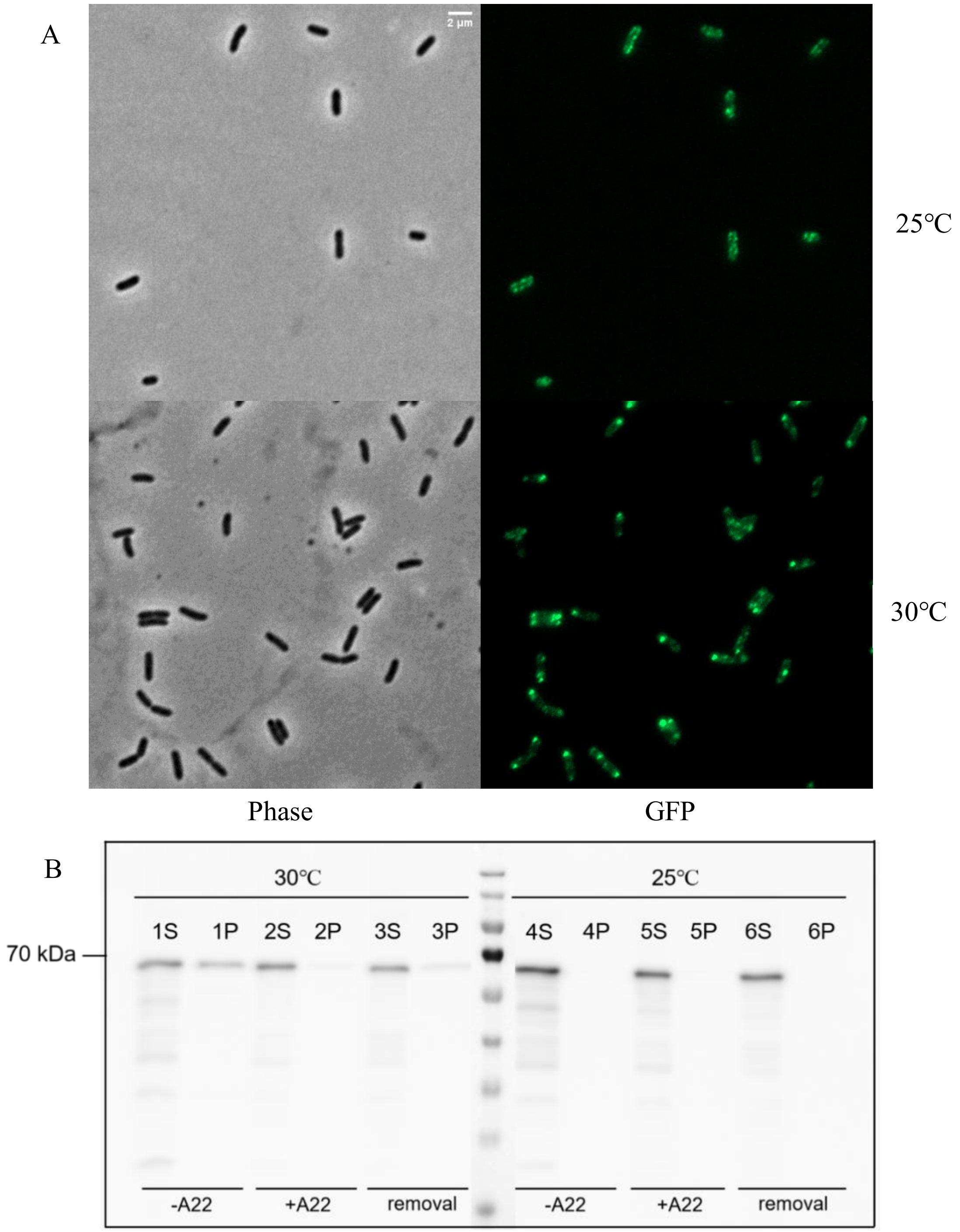

**Supplementary Figure 1.** (A) Localization of MreB-GFP in *X. citri mreB-gfp* grown at 25°C and 30°C. Phase contrast (left) and GFP (right) images were taken with the appropriate filter sets. (B)Westernblot using anti-GFP antibody to detect MreB-GFP. Supernatant (S) is the soluble fraction, Pellet (P) is insoluble material after urea extraction. Samples 1, 4: 0 µg/mL A22. 2, 5: 20 µg/mL

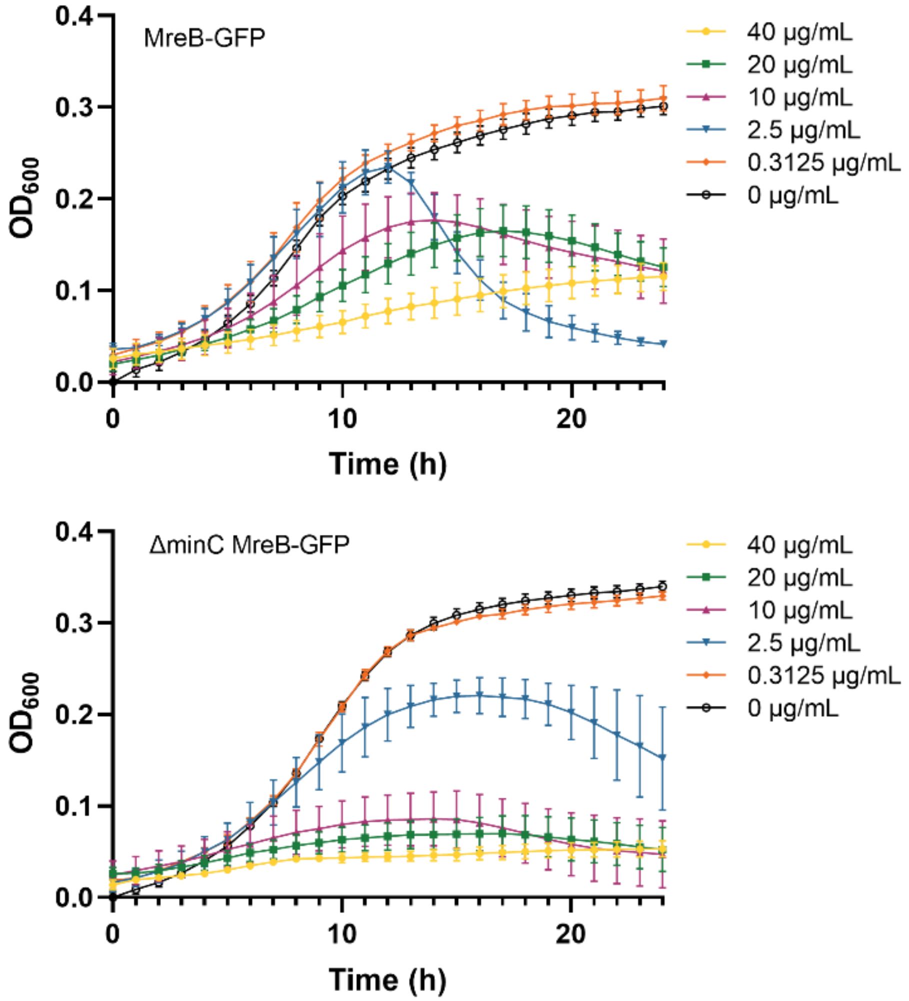

A22. 3, 6: A22 removal group. **Supplementary Figure 2.** Growth curves of *X. citri* cells in the presence of A22 at different concentrations.

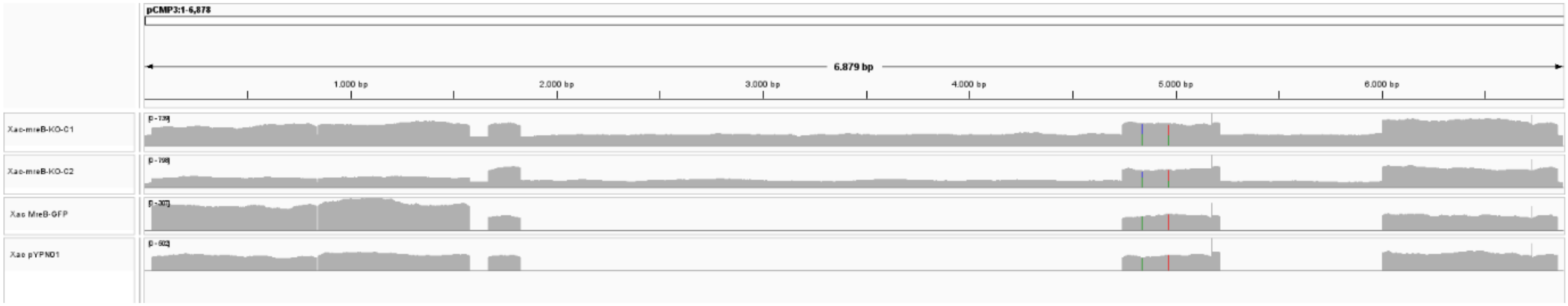

**Supplementary Figure 3.** The full suicide plasmid backbone is present in Xac-mreB-KO-C1/C2 , consistent with single-crossover integration rather than a resolved double-crossover deletion. Higher coverage regions in Xac-mreB-KO-C1/C2 are also covered in Xac MreB-GFP/Xac pYPN01 indicating these regions are present in the Xanthomonas genome.

Alignment of sequencing reads revealed the presence of the suicide plasmid in Xac-mreB-KO-C1/C2 Fig. S3 as well as some genomic regions overlapping with sections of the plasmid, as indicated by coverage of these regions in Xac MreB-GFP/Xac pYPN01 and increased coverage in sample Xac-mreB-KO-C1/C2. The non-genomic plasmid regions were also assembled into separate contigs in de novo genome assemblies created from Xac-mreB-KO-C1/C2 sequencing data.

The presence of the WT MreB and MreB-GFP encoding regions in all strains was verified by aligning sequencing reads to these regions. Reads bridging the ORF and surrounding regions indicated the presence of both ORFs within the genome Fig. S4. The presence of the WT MreB was further verified through PCR amplification of the ORF and its surrounding region with WT bands being present in all strains. Overall, this indicates Xac-mreB-KO-C1/C2 are not successful deletion mutants.

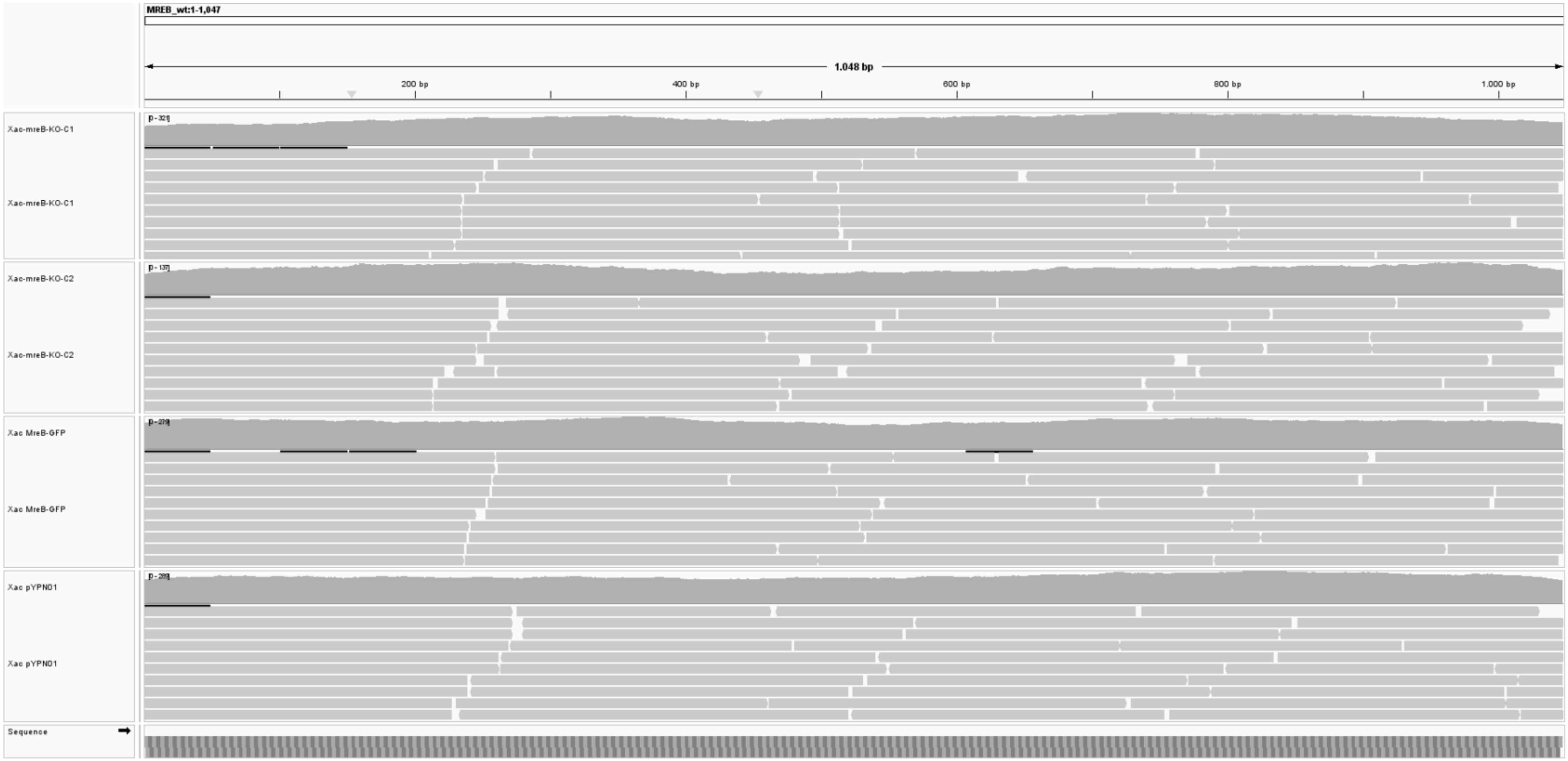

**Supplementary Figure 4.** MreB and the MreB-GFP fusion construct are present in all strains as evidenced by continuous coverage of the ORFs and their surrounding regions.

#### 3 References

Jones, L. J. F., Carballido-López, R., Errington, J. (2001). Control of Cell Shape in Bacteria. *Cell, 104*(6), 913-922. https://doi.org/10.1016/S0092-8674(01)00287-2

Khetrapal, V., Mehershahi, K., Rafee, S., Chen, S., Lim, C. L., Chen, S. L. (2015). A set of powerful negative selection systems for unmodified Enterobacteriaceae. *Nucleic Acids Research*, *43*(13), e83-e83. https://doi.org/10.1093/nar/gkv248

Lacerda, L. A., Cavalca, L. B., Martins, P. M. M., Govone, J. S., Bacci, M., Ferreira, H. (2017). Protein depletion using the arabinose promoter in Xanthomonas citri subsp. citri. *Plasmid*, *90*, 44-52. https://doi.org/https://doi.org/10.1016/j.plasmid.2017.03.005

Lorenzoni, A. S. G., Dantas, G. C., Bergsma, T., Ferreira, H., Scheffers, D.-J. (2017). Xanthomonas citri MinC Oscillates from Pole to Pole to Ensure Proper Cell Division and Shape. *Frontiers in Microbiology*, *8*, 1352. https://doi.org/10.3389/fmicb.2017.01352

Ma, W., Wang, X., Mao, Y., Wang, Z., Chen, T., Zhao, X. (2015). Development of a markerless gene replacement system in Corynebacterium glutamicum using upp as a counter-selection marker. *Biotechnol Lett*, *37*(3), 609-617. https://doi.org/10.1007/s10529-014-1718-8

Wang, T., Li, Y., Li, J., Zhang, D., Cai, N., Zhao, G., Ma, H., Shang, C., Ma, Q., Xu, Q., Chen, N. (2019). An update of the suicide plasmid-mediated genome editing system in Corynebacterium glutamicum. *Microbial Biotechnology*, *12*(5), 907-919. https://doi.org/https://doi.org/10.1111/1751-7915.13444

Zhang, Y., Buchholz, F., Muyrers, J. P., Stewart, A. F. (1998). A new logic for DNA engineering using recombination in Escherichia coli. *Nat Genet*, *20*(2), 123-128. https://doi.org/10.1038/2417
